# On Non-Kolmogorov turbulence in blood flow and its possible role in mechanobiological stimulation

**DOI:** 10.1101/2022.07.09.499422

**Authors:** Khalid M. Saqr, Iham F. Zidane

## Abstract

The study of turbulence in physiologic blood flow is important due to its strong relevance to endothelial mechanobiology and vascular disease. Recently, Saqr et al *(Sci Rep 10, 15492, 2020)* discovered non-Kolmogorov turbulence in physiologic blood flow *in vivo*, traced its origins to the Navier-Stokes equation and demonstrated some of its properties using chaos and hydrodynamic-stability theories. The present work extends these findings and investigates some inherent characteristics of non-Kolmogorov turbulence in monoharmonic and multiharmonic pulsatile flow under ideal physiologic conditions. The purpose of this work is to propose a conjecture for the origins for picoNewton forces that are known to regulate endothelial cells’ functions. The new conjecture relates these forces to physiologic momentum-viscous interactions in the near-wall region of the flow. Here, we used high-resolution large eddy simulation (HRLES) to study pulsatile incompressible flow in a straight pipe of *L*/*D =* 20. The simulations presented Newtonian and Carreau-Yasuda fluid flows, at Reynolds number of 256 and 228, respectively, each represented by one, two and three boundary harmonics. Comparison was established based on maintaining constant time-averaged mass flow rate in all simulations. First, we report the effect of primary harmonics on the global power budget using primitive variables in phase space. Second, we describe the non-Kolmogorov turbulence in frequency domain. Third, we investigate the near-wall coherent structures in time, space and frequency domains. Finally, we propose a new conjecture for the role of turbulence in endothelial cells’ mechanobiology. The proposed conjecture correlates near-wall turbulence to a force field of picoNewton scale, suggesting possible relevance to endothelial cells mechanobiology.

## 1. INTRODUCTION

The heart, as a positive displacement pump, drives arterial flow by a multiharmonic pressure waveform that is known to correlate with physiologic and pathologic processes and conditions ^1-4^. Downstream from the heart, as blood flows through the anatomically sophisticated arterial network^5^, it exhibits complex hemodynamic features such as coherent structures^6^, helical flow *(i.e. swirl)*^*7,8*^, and wave reflection and attentuation^9^.

The circle of Willis represents one of the most complex hemodynamic environments^10^ with its highly three-dimensional and intricately branching morphology^11,12^. Studies addressing the hemodynamics of the circle of Willis have been primarily motivated by the need to understand its role in neurovascular disease conditions such as intracranial aneurysm(IA)^13^ and carotid stenosis (CS)^14,15^. Intracranial hemodynamics is a key element in the disease model of neurovascular conditions and is known to be of influential role in degenerative vascular mechanisms^16,17^. The mechanobiological paradigm of such disease model is based on the endothelial mechanosensory functions^18-20^ that regulates inflammatory and atherogenic mechanisms^21,22^.

While *in vivo* measurement techniques reveal important information about vascular hemodynamics, they lack sufficient spatiotemporal resolution required to explain the biologically relevant flow characteristics. Computational Fluid Dynamics (CFD) offers an efficient framework to simulate vascular hemodynamics *in silico* by solving the notoriously difficult Navier-Stokes equation^23-25^. Similar with other CFD application areas, numerous *simplifications* are often adopted in vascular hemodynamics to solve the latter equation successfully and efficiently.

In a recent review and meta-analysis of 1733 published studies, Saqr et al^26^ showed that over 60%, 90% and 95% of IA CFD studies assumed steady, Newtonian or laminar flow, respectively. These assumptions challenge our understanding of the intracranial hemodynamic environment. In addition, the major independent flow field variable used to characterize vascular hemodynamics is wall shear stress (WSS)^27,28^. As represented in the general form of Navier-Stokes equation, WSS is a scalar-tensor field^29^ and its magnitude is calculated as the product of the symmetric elements of the stress tensor and viscosity. Therefore, WSS is not sufficient to parametrize vascular hemodynamics^26^ *in situ* hence nor to efficiently link hemodynamic characteristics with endothelial mechanosensory. That is why the correlation between WSS information and pathologic endothelial mechanotransduction remain controversial and debatable^30-33^.

A research gap can be clearly identified in the works demonstrating endothelial mechanosensory due to flow directionality^34^, regime^35^ and harmonics^36^, which cannot be correctly represented by a scalar-tensor field variable such as WSS. In fact, there are numerous studies demonstrating the role of *turbulence* in promoting proinflammatory pathways in the vascular endothelium^37-39^ which cannot find explanatory frame using WSS, as demonstrated when similar WSS values were applied under laminar flow conditions^40^. To that end, and to the best of the authors’ knowledge, there have been no attempts so far to suggest alternative fundamental hemodynamic basis for parametrizing and characterizing endothelial mechanobiology. This article is another step in the way of proposing a future research agenda in this important field.

The Kolmogorov-Obukhov theory of turbulence assumes stationary, homogenous, and isotropic decay of turbulence kinetic energy. Multiharmonic pulsatile flow, by definition, is not subject to such assumptions, and in physiologic settings, particularly in arteries, it is further excluded from such assumptions due to the non-Newtonian whole blood viscosity. The term *non-Kolmogorov turbulence* is well known in metrology where it is used to describe some atmospheric turbulence regimes. There is no unified theory to describe spatiotemporal statistics of non-Kolmogorov turbulence until today. The term is borrowed here to describe the regime(s) of turbulence observed in multiharmonic pulsatile flow where anisotropy, coherent structures and intermittent events do not follow the Kolmogorov-Obukhov theory.

Recently, Saqr et al^41^ showed that physiologic blood flow is by definition *turbulent* and explained how its regime of *turbulence* go beyond the Kolmogorov-Obukhov theory of homogenous isotropic *turbulence*. A direct implication of such work is the necessity of developing new hemodynamic variables that can parametrize endothelial mechanosensory with respect to the characteristics of blood turbulence. To achieve this, such rarely studied regime of turbulence should be characterized first. This work is an attempt to do so by means of high resolution Large Eddy Simulation that has recently showed excellent performance in investigating intracranial hemodynamics associated with transient ischemic attack^42^, carotid stenosis^43^ and cerebral aneurysm^44,45^. The main objective of this work was to investigate the effects of primary blood waveform harmonics on blood flow dynamics and turbulence characteristics. The secondary objective was to qualitatively describe the near-wall turbulence region and correlate it with the picoNewton forces that are related to endothelial mechanosensory.

## 2. METHODS: HIGH RESOLUTION LARGE EDDY SIMULATION

Six CFD simulations comprising a Newtonian and a non-Newtonian sets, each presenting three sets of boundary condition representing different primary harmonics, were solved using Large Eddy Simulation (LES) with the Wall-Adaptive Local Eddy-Viscosity (WALE) subgrid model on structured Cartesian grid with central-differencing spatial discretization and second order implicit time stepping, as reported previously by Rashad et al^42^. The Courant number in all reported cases was maintained below unity. Eight pulses were solved, three pulses were discarded and five were analyzed for each simulation case. Simulations were carried out on 8-core Intel Core i7 computing node with 32 GB RAM and each case produced 300 GB of data and consumed approximately 156 CPU-hours. Spatial convergence was ensured at each time step with absolute residuals below 10^−5^ for the velocity and pressure and 10^−6^ for turbulent viscosity fields. The models were implemented and solved using *OpenFOAM* v2106^46^.

### 2.1. Governing equations

The present computational work solves filtered Navier-Stokes equation that can be expressed as:

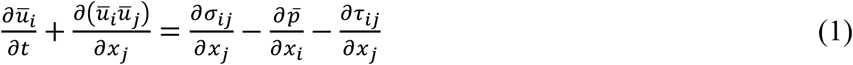

where *σ*_*ij*_ is the viscous stress tensor defined as:

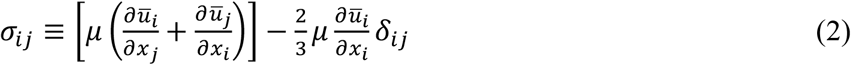

and *τ*_*ij*_ is the subgrid stress tensor defined for incompressible flow as:

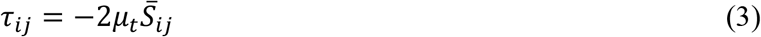

where 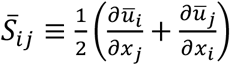 and *μ*_*t*_ is calculated from the Wall-Adaptive Local Eddy Viscosity model^47^ that is expressed as:

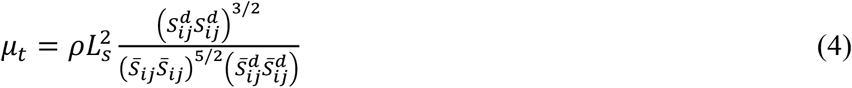

where *L*_*s*_ and 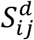 are the subgrid filter length and the traceless symmetric part of the square of the velocity gradient tensor, respectively, expressed as:

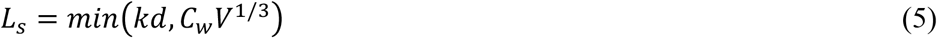

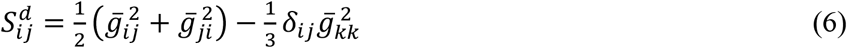

where 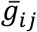 is the velocity gradient tensor 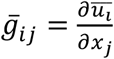 and *δ*_*ij*_ is the Cartesian Kronecker operator. The value of WALE constant *C*_*w*_ was taken as 0.325. The WALE model was chosen as it offers significant advantages in modeling wall-bounded turbulence compared to other mainstream SGS models^48^.

The Newtonian models had a constant viscosity of 0.0035 pa.s while the non-Newtonian models had effective viscosity calculated via the Carreau-Yasuda model^49^ expressed as:

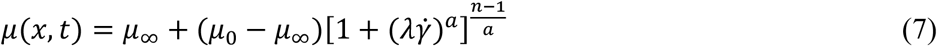

where *μ*_∞_=0.0022 Pa.s, *μ*_0_=0.022 Pa.s, *λ*=0.11 s, *a*=0.644, n=0.392 according to Gijsen et al^50^ and 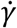 is the local instantaneous shear rate magnitude.

It is noteworthy to mention here that the basis of comparison between the Newtonian and non-Newtonian models is the global value of the mean shear-dependent viscosity 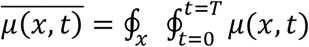, as shown in a recent study by Saqr^51^. In all the six cases presented here, such value was found to 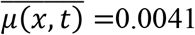 Pa.s, which is different than the Newtonian viscosity by 17.1%. In experimental measurements^52-54^, similar, and sometimes higher, differences between Newtonian and non-Newtonian fluids can be generally accepted for comparing both viscous and inviscid invariants of the velocity field. Therefore, the authors believe that it is an accepted variation margin in their computational work and the comparison is therefore representative of the empirical reality.

### 2.2. Boundary conditions

In the carotid artery, where 95% of blood flow harmonic frequencies are below 12 Hz, the top three harmonics, in terms of amplitude, contain frequencies less than 6 Hz ^55^. Therefore, the harmonics used in the present work were selected in the range of 1-4 Hz. A straight tube with *L*/*D* = 20 was designated to represent ideal flow in the cervical segment (C1) of the internal carotid artery. Boundary conditions were set as a velocity inlet time series and constant pressure on the outlet. The velocity time series was expressed as:

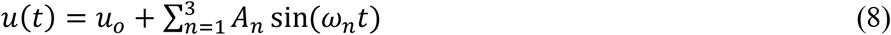

where *n* refers to the waveform harmonic index indicating amplitude *A*_*n*_ and frequency *f*_*n*_ *= ω*_*n*_/2*π*. The assumption of parabolic velocity profile was skipped in the present work to allow the flow to form coherent structures in the near-wall region as it develops along the tube to mimic the structures formed due to morphological features such as tortuosity and bifurcations^42^. The cases and corresponding boundary conditions are detailed in table 1 and the waveform harmonics are shown in figure 1.

**Table 1.** Details of the boundary conditions of the six LES cases as represented by equation (8). N and CY stand for Newtonian and Carreau-Yasuda models. The steady velocity component was fixed at: *u*_*o*_ *=* 0.175*m*/*s*

| Case | $A_n(\text{m/s}), f_n(\text{Hz})$ | | |
| --- | --- | --- | --- |
| | $n = 1$ | $n = 2$ | $n = 3$ |
| **1N** | 0.1, 1 | - | - |
| **1CY** | 0.1, 1 | - | - |
| **2N** | 0.1, 1 | 0.05, 2 | - |
| **2CY** | 0.1, 1 | 0.05, 2 | - |
| **3N** | 0.1, 1 | 0.05, 2 | 0.025, 4 |
| **3CY** | 0.1, 1 | 0.05, 2 | 0.025, 4 |

**Figure 1.**
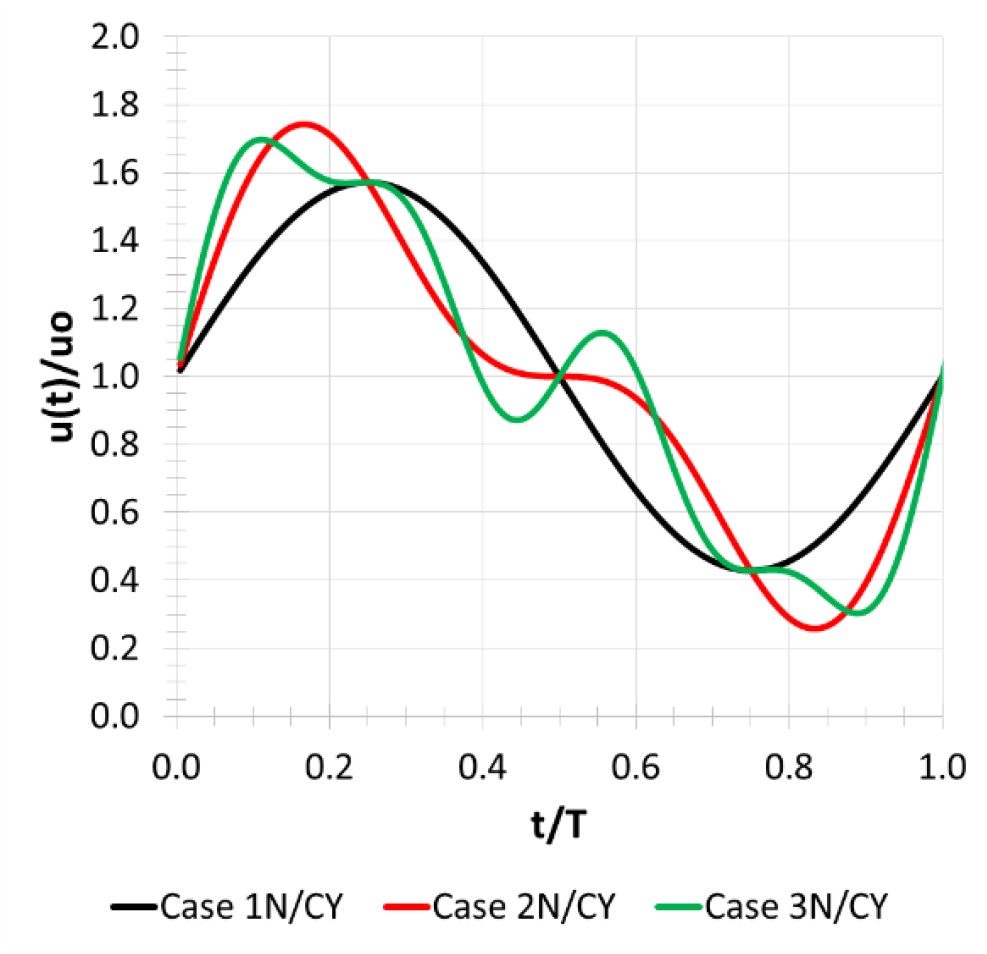
Boundary conditions waveforms presenting one harmonic (case 1N/1CY), two harmonics (case 2N/2CY) and three harmonics (case 3N/3CY) where the X and Y axis represent dimensionless time and velocity values for one pulse.

The mean Reynolds number of the Newtonian and Carreau-Yasuda models was 265 and 228, respectively. Non-Newtonian mean Reynolds number was calculated using the generalized Metzner-Reed correlation. The values of the harmonic amplitude and frequency were determined such that the time-averaged flow rate at the boundary for all cases is kept constant at 0.1741±5% *m*^3^/*s* to establish inertial basis for comparing the momentum-viscous interactions, as shown later in the results and discussion.

### 2.3. Computational grid and LES quality assessment

In pulsatile blood flow, turbulence is generated due to intermittency resulting from the inherited nonlinearity of the primary harmonics as shown analytically^41^ and experimentally^56,57^. We have conducted a grid sensitivity test to estimate the required mesh resolution and to demonstrate the capabilities of our LES solution strategy.

The criteria proposed by Pope^58^ was adopted as a measure for LES quality assessment. Such criteria defines *M*(*x*_*i*_, *t*) *= K*_*res*_/*K*_*tot*_ as a scalar field that represents the ratio of resolved (*K*_*res*_) to total (*K*_*tot*_ *= K*_*res*_ + *K*_*SGS*_) turbulence kinetic energy in any LES solution, where subscripts *res, SGS, tot* represent resolved, sub-grid scale and total turbulence kinetic energy, respectively. Pope showed that if such scalar is higher than 0.85, the LES solution could be deemed to represent the closest solution to a Direct Numerical Simulation (DNS).

Three computational grids (i.e. meshes) with cell count of 0.1, 0.5 and 1 million were created and solved with LES under five cycles of monoharmonic pulsatile flow conditions corresponding to *Re*_*m*_ = 265 and 454, while 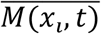 values were calculated in runtime using a C# User Defined Function (UDF) as previously explained by the authors^42^. Figure 2 shows the results of the grid sensitivity analysis. The minimum and maximum values of of 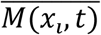 correspond to peak systole and minimum diastole conditions. It is shown that mesh2 and mesh3 achieve Pope’s criteria for both values of *Re*_*m*_ with resolution of 0.5 and 1 million cells, respectively. The computing time for mesh3 was 2.1 times higher than such of mesh2. Therefore, mesh2 was selected to conduct the LES analysis. This grid had maximum *y*^+^ value of 0.14 during the solution of all cases.

**Figure 2.**
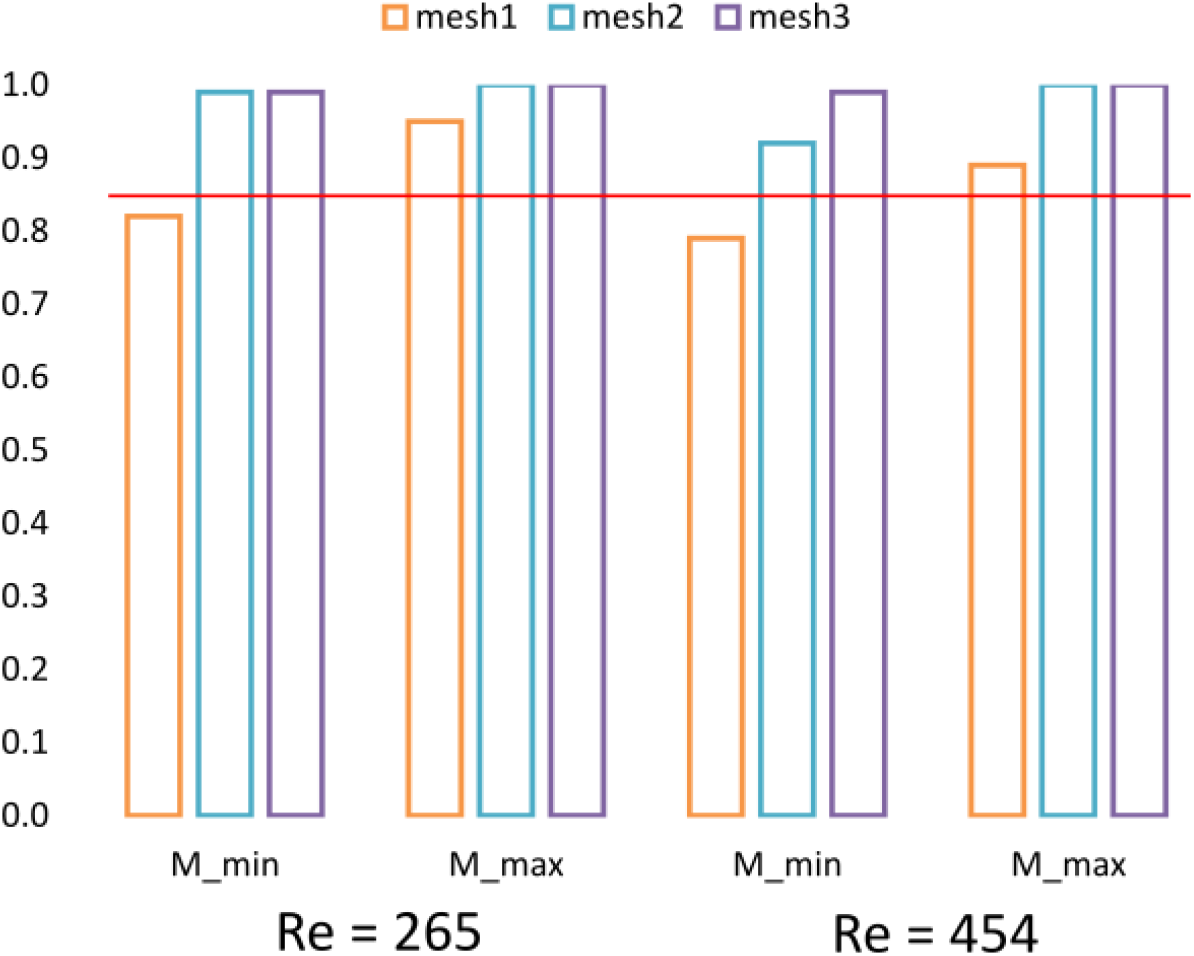
The minimum and maximum values of 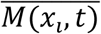 on y-axis for computational grids of 0.1, 0.5 and 1 million mesh cells for mesh1, mesh2 and mesh3, respectively.

## 3. RESULTS AND DISCUSSION

Three sets of results were analyzed from each simulation. Two datasets were obtained at two vertex stations, the first (C) is located on the centreline of the computational domain at an axial distance *L*/*D =* 10 from the inlet plane. The second station (NW) is at the same axial distance, however, located 10*μm* from the wall. The third dataset was obtained as isovolumetric maps at peak systolic and minimum diastolic instances. The frequency analysis was initially conducted with a 1 *kHz* sampling rate and then resampled to 40 *Hz* to cut off energy levels below 10^−16^ *m*^2^/*s*^2^. The resampled datasets represent the frequencies covered 99.1% of the mass transfer process in all simulations and therefore they were used to present the results.

### 3.1. Hydrodynamic phase shift

In pulsatile flow, there is an inherent phase shift between velocity and pressure^57,59^ due to the inertial effects that can be justified by comparing the terms *ρV*. ∇*V* and ∇*p* under non-zero acceleration conditions. Considerable efforts have been done to investigate such phenomena^60-62^, however, most of the works investigated monoharmonic flows and Newtonian fluids.

In monoharmonic flow, the phase shift between pressure and velocity is uniform and constant, following an elliptic profile^63^. This is shown in figure 3 (top row). The phase shift is different from the center to the near-wall flow, due to the viscous effects at the wall. When a second harmonic is introduced to the flow, as shown in figure 3 (middle row), the phase shift changes within the cycle forming the non-uniform kidney-shaped map shown in the figure. In the near-wall region, the non-uniformity is more evident. Adding a third harmonic brings irregularity to the phase map, as shown in figure 3 (bottom row). In this case, the phase shift varies in more complex way creating three distinct features in the phase map that is also more complex near the wall.

**Figure 3.**
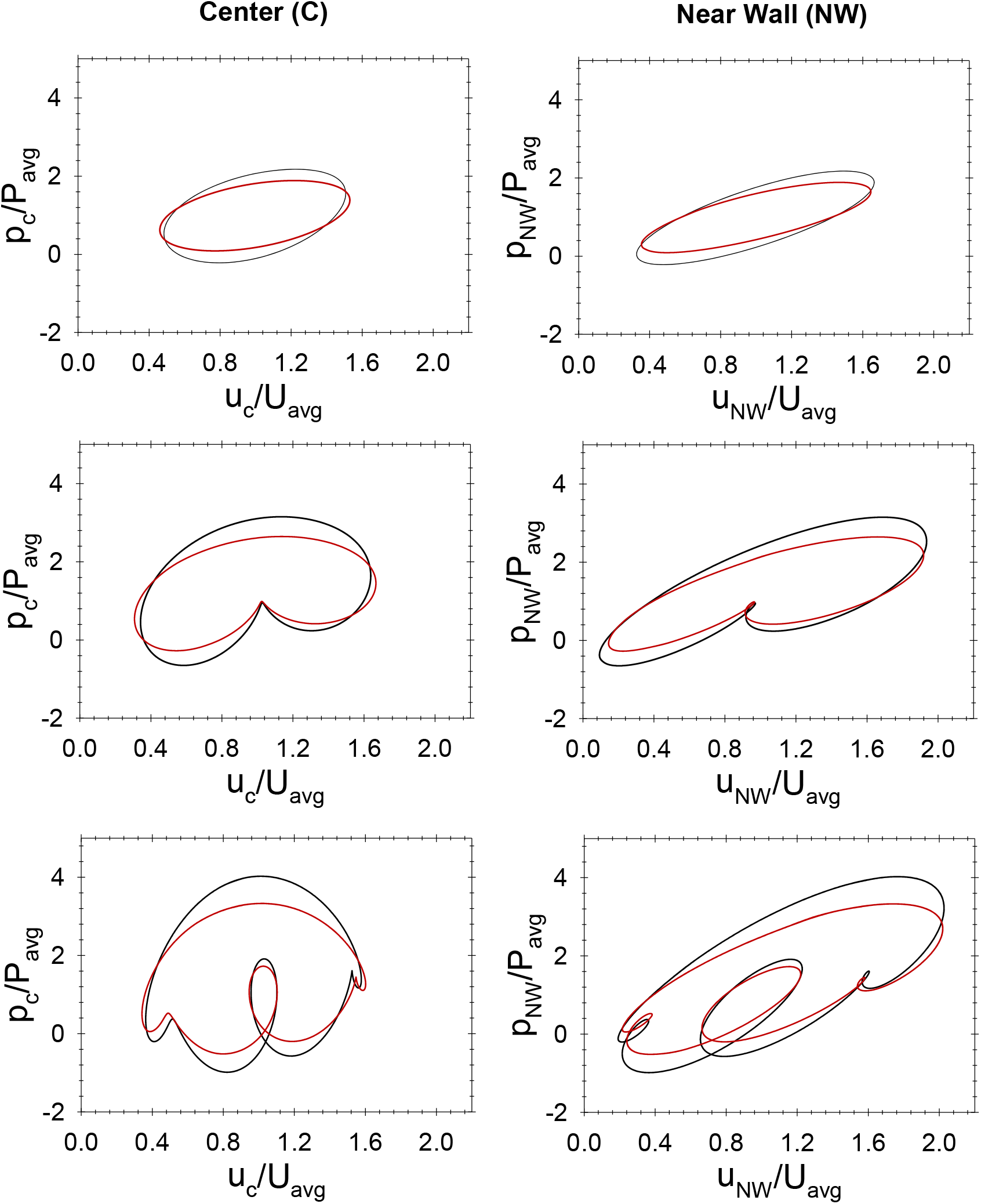
Non-dimensional phase diagrams of pulsatile flow showing the effect of harmonics (one, two and three harmonics depicted in the top, middle and bottom rows, respectively). Black and red lines indicate Newtonian (N) and Carreau-Yasuda (CY) fluid models, respectively. Spatial non-similarity is shown by comparing the phase diagrams obtained at a center point (left column) and a near-wall pint 10 *μm* from the wall (right column). Average values are calculated over one cycle for each simulation.

The phase shift variation indicates the nonlinear effects of adding a second, then a third, harmonic to the flow. The superposition of harmonics complicates the competition between inertia and pressure in space and time. The consequences of this phenomena on the biologically relevant flow dynamics could be fundamental. The area inside the phase map represents the local dimensionless work exerted in the system. Despite that the mass flow rate per cycle was maintained constant in all simulations, the Carreau-Yasuda models (YC) had 7-12% less work than the Newtonian models (N). In all cases, nevertheless, the system exerted more work in the near-wall (NW) region than in the center region (C), which can be explained by the significance of the viscous term *μ*(∇^2^*V*) in the near-wall region.

### 3.2. Turbulence Kinetic Energy and Vortex Breakdown

The turbulence kinetic energy (TKE) cascade was analyzed in the center and near-wall stations for the six models solved here. It was only possible to present the TKE cascade in frequency domain. This is because Taylor’s frozen turbulence hypothesis does not hold and there is no way of correlating time and length scales. Figure 4 shows the TKE cascade plots for monoharmonic flow (top row) and for pulsatile flow with two harmonics (middle row) and three harmonics (bottom row) for Newtonian and Carreau-Yasuda fluids.

**Figure 4.**
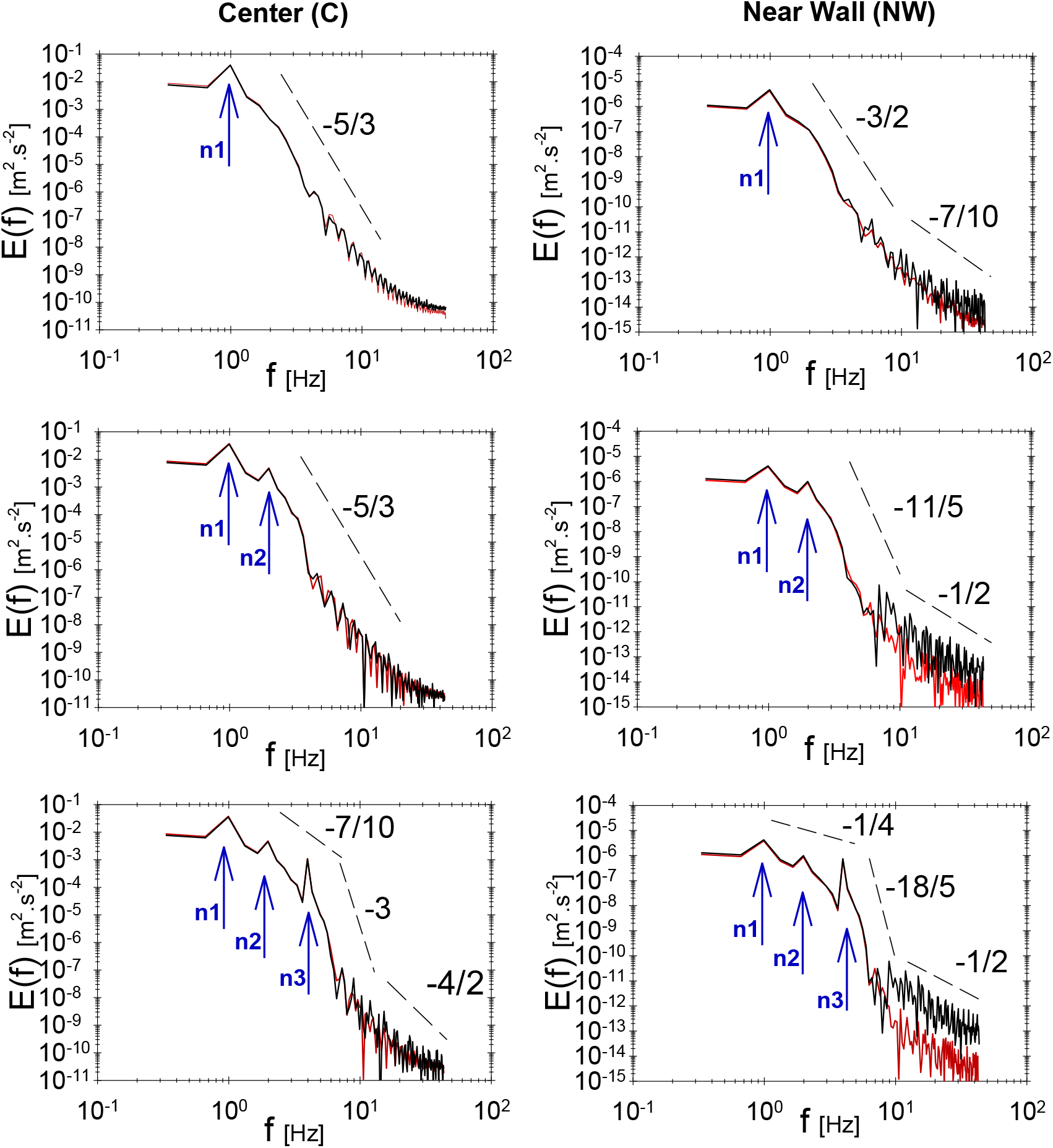
Turbulence Kinetic Energy E(f) cascade in frequency domain (f) at the center point (left column) and 10 *μm* near-wall (right column). The depicted cascades present flows with one harmonic (n1), two harmonics (n1,n2) and three harmonics (n1,n2,n3) in the first, second and third rows, respectively. Black and red lines represent Newtonian and non-Newtonian fluids, respectively.

#### 3.2.1 Primary harmonics traces in TKE cascade

The primary harmonics (*n*1 − *n*3) can be traced in the TKE cascade plots as spikes at the corresponding frequency values, as depicted in figure 4. The primary harmonics represent the source of kinetic energy that is provided to the flow while the secondary harmonics, depicted at frequencies other than the primary frequencies, represent the flow scales and structures. Comparing TKE cascade in the center and near-wall, it is observed that primary harmonics have 4 orders of magnitude difference between the two locations.

#### 3.2.2 TKE Cascade and scaling

In order to illustrate the features of the non-Kolmogorov turbulence reported here, it is vital to describe both the cascade (i.e. decay) as well as the levels (i.e. magnitude) of TKE. Since it is not possible to apply Taylor’s frozen turbulence hypothesis here, it is only possible to describe TKE in frequency domain. In monoharmonic pulsatile flow, figure 4 (top row), the TKE cascade depict a single plateau cascade with a magnitude-range from 10^−1^ *m*^2^/*s*^2^ to 10^−11^ *m*^2^/*s*^2^ at a slope corresponding to −5/3 in the center point of the flow. On the other hand, in the near-wall region, it is possible to observe two plateaus. The first has a slope corresponding to −3/2 and is bounded by the primary harmonic of 10^−5^*m*^2^/*s*^2^ and 1 Hz and 10^−12^ *m*^2^/*s*^2^ and 8 Hz in magnitude and frequency, respectively. The second plateau refers to small scale oscillations with a cascade slope corresponding to −7/10 and shows TKE oscillations ranging from 10^−12^ *m*^2^/*s*^2^ at 10 Hz to 10^−15^*m*^2^/*s*^2^ at 30 Hz in magnitude and frequency, respectively, in the near wall region.

Adding one and two harmonics to the flow, as shown in figure 4 (middle and bottom rows, respectively) changes the TKE cascade, particularly in the near-wall region. The increasing difference between center (C) and near-wall (NW) TKE cascades, in terms of cascade slope, bounds, and magnitude, further demonstrate the non-Kolmogorov features of the turbulence field. The near-wall TKE cascade is characterized by an increase in the TKE oscillations magnitude at the frequency range of 10 ≤ *f* ≤ 30.

It is also important to note that the viscous effects appear only in the near-wall region where the Carreau-Yasuda fluid demonstrate less levels of energy, by approximately one order of magnitude, than the Newtonian fluid. In a physical sense, it means that the dissipation of TKE from inertial to viscous scales is more rapid in Carreau-Yasuda fluid. In the low-frequency range 1 ≤ *f* < 8, viscous effects do not exist as the strain-rate is negligible compared to vorticity in large scales. The dominant effect of momentum transfer spans the primary harmonics range and replaces the Kolmogorov-inertial subrange commonly observed in fully developed turbulence. In the case of three harmonics, however, there is an intermediate plateau appears after the third harmonic spike (*n =* 3) and brings an abrupt four-orders of magnitude drop in energy at 4 ≤ *f* ≤ 7, as in figure 3 (bottom row). This effect is observed in the center (C) and near-wall (NW), however, with an evident quantitative difference between Newtonian and Carreau-Yasuda fluids in (NW).

#### 3.2.3 Near-wall coherent structures

In physiologic conditions, the irregularity and complex 3D morphology of the vascular network produce vortex structures in blood flow. Streamwise vortex structures are the dominant type of structures in arteries^64-66^. To synthesize streamwise vortex structures in the present ideal model, flow was admitted to the computational domain with a plug-flow profile^67^. Differential drag in the entrance region produced ring vortices of significant scales that propagates in streamwise direction and undergoes shedding and breakdown.

We used Q-criterion^68^ 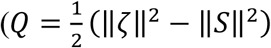, where ζ and *S* are the vorticity and strain rate tensors) to investigate the qualitative features of coherent structures and their variation according to harmonics, viscosity, and cardiac cycle. Although it is important to investigate the coherent structures quantitatively, the authors could not find an appropriate method in literature and are currently developing one. In figure 5, iso-volumes of positive Q-criterion regions (0 < *Q* ≤ 0.02) are plotted with a colour map representing vorticity magnitude. These iso-volumes represent the near-wall persisting structures that form along the flow domain due to the breakdown on the entrance ring vortex. It is obvious that the effects of local acceleration, as represented by different instances (rows, right) of the cardiac cycle, control the vorticity field magnitude. It is also clear that the viscosity (N, CY, sub-rows left) has diminishing effect on the structure and magnitude of the near-wall structures. The number of harmonics affects the vorticity magnitude at the same instance of time (*n* = 1,2,3, columns). In minimum diastole, cases 2N and 2CY exhibit near-zero vorticity field. In mid deceleration, cases 3N and 3CY have approximately 20% increase in the vorticity magnitude compared to cases 1N and 1CY.

**Figure 5.**
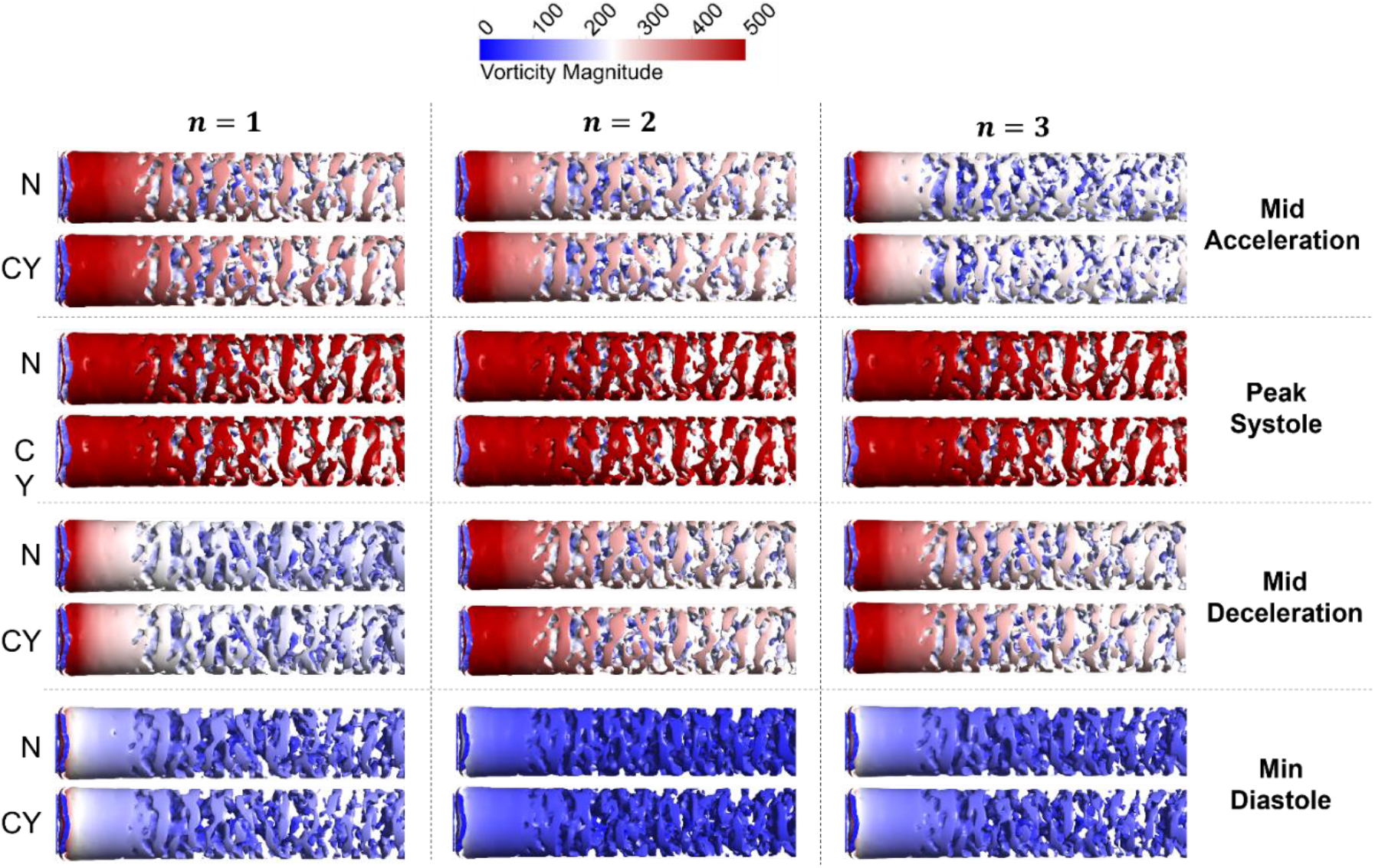
Iso-volumes of Q-criterion coloured by vorticity magnitude at (0 < *Q*≤ 0.02, 0 ≤ ‖ζ‖ ≤ 500) for different harmonics (*n*=1,2,3, columns) at different time instances of the cardiac cycle (rows, right) for Newtonian and non-Newtonian viscosity (N and CY, rows, left).

## 4. DISCUSSION

Experimental measurements carried out by Brindise and Vlachos^69^ showed that blood flow turbulence is always generated at the wall. Their PIV measurements showed that hemodynamic turbulence depends on the boundary waveform (i.e. harmonics). Feaver et al^70^ showed that the blood flow harmonics regulate endothelial inflammation, including NF-kB and downstream inflammatory phenotype. Molla et al^71^ used Large Eddy Simulation (LES) to study *transitional* pulsatile flow through a stenotic channel. They showed that TKE cascade slopes vary significantly from Kolmogorov’s scaling laws. Their work was later supported by Lancellotti et al^72^, Mancini et al^73^, Ozden et al^74^, and most recently by Saqr et al^75^.

Our results showed that boundary harmonics that the shape of blood flow waveform play a key-roles in the near-wall flow field by controlling TKE cascade and underlying vortex breakdown dynamics. The near-wall flow is characterized by non-Kolmogorov turbulence, confirming the recent findings by Saqr et al^75^. The near-wall flow field is the hemodynamic environment of endothelial cells. Such environment can be conjectured to be mainly regulated by harmonics. Low frequency harmonics inherited from cardiac waveforms, mostly associated with coherent structures, regulate mass flow and quasi-steady events in energy and vortex breakdown. High frequency harmonics, generated in the flow from vortex breakdown, regulate viscous dissipation and its associated TKE cascade rates. Persisting near-wall coherent structures communicate a *vortex force* field to the wall that is lined with endothelial cells in physiologic conditions. The association between TKE intermittency and vortex breakdown is well documented in the literature of fully developed turbulence^76^. However, there is no generalized exact mathematical definition for vortex breakdown while there is consensus that it can be generally described as sudden change in vortex structure^77^. The generation and dissipation of a vortex is associated with the change of forces on the walls from where the vortex originates and can be expressed as 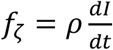 where *I* is the Lamb vector *I* = *V*×ζ that balances the solenoidal part of the convective derivative 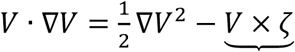. The impulse of a vortex ring of a circulation Γ = ∬_*S*_ ζ*ds* and radius R has only one streamwise component that can be expressed as *M*_*x*_ = *ρ*Γπ*R*^2^. If the time variation of the radius can be neglected with respect to the circulation variation, the vortex force becomes a function of the rate of change of circulation, as 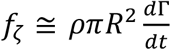. This vortex force is resulting from the local acceleration field 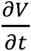 that is inherent in physiologic flows. In which, the vortex force produces a continuous hammering^78^ localized onto the region of the tissues where the vortex develops. Such hammering can rationally be correlated with the cell-scale response to picoNewton forces^79^.

While it is important to explore the coherent structures in vascular flows, it is important to realize that such exploration is still an open problem until today^80^. Vascular flows produce surprisingly complex patterns in space and time such that fluid dynamicists have yet to find the perfect variables to study them. The use of Q-Criterion expresses the mainstream direction in the community where coherent structures are identified as persistent vortex rings and formations^81,82^. However, attempts to provide better ways of visualizing coherent structures have been recently reported. Calò et al^83^ proposed, for the first time, a complex network based-approach to capture the large-scale coherent structures from 4D MRI imaging data of ascending aortic aneurysm (AAo). They demonstrated how their complex networks approach could be useful to quantitative *in vivo* hemodynamic risk assessment of such fatal condition. It should also be noted that the integration of CFD-based hemodynamic markers is essential in developing next-generation personalized management systems and digital twins for cardiovascular patients.

## 5. CONCLUSION

This study compliments a growing body of literature by the author and other research teams exploring vascular hemodynamics beyond the WSS paradigm. Despite the invariance of time-averaged flow rate, harmonics was evidently shown to regulate both viscous and inviscid flow field variables of both Newtonian and non-Newtonian fluids. By increasing the number of harmonics, the power budget of the flow increased and a noticeable influence of viscosity on TKE cascade rates is found. The turbulence regime of the flow demonstrates non-Kolmogorov features with observable differences between near-wall and main flows particularly in high-frequency regime. The corresponding coherent structures were found to have vorticity magnitude regulated by harmonics. The authors find it reasonable to conclude that the resulting vortex force field, that corresponds to TKE energy levels within 10^−11^ − 10^−15^*m*^2^. *s*^−2^, regulates endothelial cells mechano-functions and represent a rational explanation of the cell-scale mechano-response to piconewton forces that have been previously reported in literature.

## Author Contributions

**KS** conceived the study, designed the methodology and planned the parametric study. **IZ** conducted the CFD setup and simulations. **KS & IZ** conducted the verification and LES grid quality assessment. **KS & IZ** interpreted the results and conducted the post-processing. **KS** wrote the manuscript.

## FUNDING

None.

## DATA AVAILABILITY

Data would be made available upon request from the authors.

## Notes

### Competing Interest Statement

The authors have declared no competing interest.

